# RppH can faithfully replace TAP to allow cloning of 5’-triphosphate carrying small RNAs

**DOI:** 10.1101/283077

**Authors:** Miguel Vasconcelos Almeida, António Miguel de Jesus Domingues, Hanna Lukas, Maria Mendez-Lago, René F. Ketting

**Affiliations:** Biology of Non-coding RNA Group, Institute of Molecular Biology, 55128 Mainz, Germany; Genomics Core Facility, Institute of Molecular Biology, 55128 Mainz, Germany

## Abstract

RNA interference was first described in the nematode *Caenorhabditis elegans*. Ever since, several new endogenous small RNA pathways have been described and characterized to different degrees. Much like plants, but unlike *Drosophila* and mammals, worms have RNA-dependent RNA Polymerases (RdRPs) that directly synthesize small RNAs using other transcripts as a template. The very prominent secondary small interfering RNAs, also called 22G-RNAs, produced by the RdRPs RRF-1 and EGO-1 in *C. elegans*, maintain the 5’ triphosphate group, stemming from RdRP activity, also after loading into an Argonaute protein. This creates a technical issue, since 5’PPP groups decrease cloning efficiency for small RNA sequencing. To increase cloning efficiency of these small RNA species, a common practice in the field is the treatment of RNA samples, prior to library preparation, with Tobacco Acid pyrophosphatase (TAP). Recently, TAP production and supply was discontinued, so an alternative must be devised. We turned to RNA 5’ pyrophosphohydrolase (RppH), a commercially available pyrophosphatase isolated from *E. coli*. Here we directly compare TAP and RppH in their use for small RNA library preparation. We show that RppH-treated samples faithfully recapitulate TAP-treated samples. Specifically, there is enrichment for 22G-RNAs and mapped small RNA reads show no small RNA transcriptome-wide differences between RppH and TAP treatment. We propose that RppH can be used as a small RNA pyrophosphatase to enrich for triphosphorylated small RNA species and show that RppH- and TAP-derived datasets can be used in direct comparison.

## Introduction

RNA interference (RNAi), the process whereby small RNAs and their cognate Argonaute proteins regulate gene expression, was initially discovered in the nematode *Caenorhabditis elegans* (Fire et al., 1998). Interestingly, in nematodes several endogenous small RNA pathways and 27 genomically-encoded Argonautes are present (Billi et al., 2014). Two classes of endogenous small RNAs are transcribed by RNA-dependent RNA Polymerases (RdRPs), the 22G- and 26G-RNAs. The 22G-RNAs exist in large numbers and, consensually, have 22 nucleotides with a 5’ bias for guanosine (Billi et al., 2014). RdRP transcription leaves a triphosphate group at the 5’ end of the small RNA and consequently, mature 22G-RNAs still have this RdRP signature at their 5’ end (Billi et al., 2014). Mature 26G-RNAs do not have a 5’-triphosphate group, but it is currently unclear how this is achieved.

The 5’ triphosphate group decreases clonability of 22G-RNAs in standard small RNA library preparations. A common practice in the field to improve the yield of 22G-RNA reads in deep-sequencing experiments is to treat samples, prior to library preparation, with Tobacco Acid Pyrophosphatase (TAP) (Gu et al., 2009). TAP hydrolyzes several types of pyrophosphate bonds, including the eukaryotic 5’ Cap, but it does not hydrolyze polymerized RNA and DNA (Shinshi et al., 1976). TAP can also hydrolyze pyrophosphate groups from small RNAs, for example 22G-RNAs, increasing the likelihood of cloning 22G-RNAs during library preparation. This type of treatment is essential to address aspects of 22G-RNA biology. For example, 22G-RNAs provide transcriptome-wide self- and non-self recognition platforms that silence foreign gene expression or license bona-fide gene expression, respectively (Ashe et al., 2012; Bagijn et al., 2012; Claycomb et al., 2009; Lee et al., 2012; Luteijn et al., 2012; Seth et al., 2013; Shirayama et al., 2012; Wedeles et al., 2013; van Wolfswinkel et al., 2009). Also, several studies focused on understanding the dynamics of inheritance and establishment of silencing by these small RNA species (Houri-Ze’evi et al., 2016; Sapetschnig et al., 2015).

Recently, TAP production was discontinued, so we sought a commercially available alternative. We chose *E. coli* RNA 5’ Pyrophosphohydrolase (RppH). RppH primes mRNA degradation in *E. coli* by converting the 5’ triphosphate into a 5’ monophosphate (Deana et al., 2008). In practical terms, RppH is an enzyme analogous of TAP, allowing for a direct replacement of TAP in the established protocols, and it is available commercially at a comparable price. Here, we treated small RNAs from *C. elegans* with TAP and RppH and compared their performance in regard to 22G-RNA enrichment. We show that 22G-RNAs were similarly enriched in both treatments and there were no global differences in mappability to the *C. elegans* genome. Furthermore, RppH-treatment retrieves 22G-RNA sequences targeting known 22G-RNA regulated genes to an extent comparable to TAP-treated samples. Therefore, RppH is a viable and affordable alternative to TAP for 22G-RNA enrichment and TAP- and RppH-treated libraries can be compared directly.

## Materials and methods

### Worm culture, RNA extraction and enrichment for small RNAs

*Caenorhabditis elegans* populations were grown according to standard conditions (Brenner, 1974). Wild-type N2 worms were grown at 20°C on nematode growth medium (NGM) plates seeded with *E. coli* OP50. For synchronization, gravid adult worm populations were bleached and eggs were allowed to hatch in an overnight incubation at 20°C, in M9 buffer. Synchronized L1 larvae were plated the next day on OP50-seeded plates and allowed to grow at 20°C for 60-63 hours until adulthood. Gravid adults were washed off plate with M9 buffer and lysed in 250 μL of Worm Lysis Buffer (0.2M NaCl, 0.1M Tris pH 8.5, 50 mM EDTA, 0.5% SDS) with 150 μg Proteinase K (Sigma-Aldrich #P2308) for at least 30 minutes at 65°C. 750 μL of Trizol LS (Life Technologies, 10296-028) were added per sample and extraction followed manufacturer’s instructions, with the exception that phase-lock tubes were used to facilitate phase-separation. Total RNA extraction was followed by TURBO DNase treatment (Life Technologies #AM1907) according to manufacturer’s instructions. Small RNAs were isolated from DNase-treated total RNA using MirVana (Life Technologies #AM1561).

### RppH and TAP treatments and library preparation

#### Pre-treatment

A single RNA sample was divided into 6 technical replicates. Three of these technical replicates were treated with Tobacco Acid Phosphatase (TAP, Epicentre) and the other three, with RNA 5’Pyrophosphohydrolase (RppH, New England Biolabs, #M0356S). For TAP-treated samples, 1 μg of total RNA was incubated for 2 hours at 37°C, with 5 units of TAP enzyme and 10X TAP buffer. For RppH treated samples, 1μg of RNA was incubated for 1 hour at 37°C, with 5 units of RppH enzyme and 10X NEB Buffer 2. To stop the RppH treatment, 500mM EDTA was added and samples were incubated for 5 minutes at 65°C.

#### Library preparation

Small RNAs (16-30nt) were enriched by performing size selection of the RNA prior to library preparation. RNA samples were run on a 15% TBE-Urea polyacrylamide gel (BioRad), and purified with sodium chloride/isopropanol precipitation. Library preparation was performed using the NEBNext Multiplex Small RNA Library Prep Set for Illumina (New England BioLabs) as recommended by the manufacturer (protocol version v2.0 8/13), but replacing the RNA ligation adapters with adapters with 4 additional random bases at their 3’ ends to allow identification of PCR clonal artefacts (synthesized by Bioo Scientific). Fourteen PCR cycles were used for library amplification The PCR-amplified cDNA was purified using AMPure XP beads (Beckman Coulter). Size selection of the small RNA library was done on LabChip XT instrument (Perkin Elmer) using DNA 300 assay kit extracting the 135-170 bp fraction.

#### Next generation sequencing

The 135-170 bp library fractions of the RppH and TAP-treated libraries were pooled in equal molar ratio. The resulting 4 nM pool was denatured and diluted to 9 pM with 5% PhiX spike-in and sequenced as single-read on MiSeq (Illumina) for 68 cycles. Libraries for untreated samples, originating from a different biological replicate, were also prepared using the protocol above, but were sequenced as single-read on HiSeq 2500 (Illumina). Untreated samples were originally used in (Almeida et al., 2018).

### Bioinformatics analysis

#### Read processing and mapping

The quality of raw sequenced reads was accessed with FastQC, illumina adatpers were then removed with cutadapt (-O 8 -m 26 -M 38), reads with low-quality calls were filtered out with fastq_quality_filter (-q 20 -p 100 -Q 33). Using information from unique molecule identifiers (UMIs) added during library preparation, reads with the same sequence (including UMIs) were collapsed to removed putative PCR duplicates using a custom script. Prior to mapping, UMIs were trimmed (seqtk trimfq) and library quality re-assessed with FastQC. Reads were aligned against the *C. elegans* genome assembly WBcel235 with bowtie v0.12.8 (--tryhard --best --strata --chunkmbs 256 -p 8 -v 1 -M 1). For the analysis, only reads mapping to a single genomic location were used. Except where indicated, reads used for comparison are those whose sequence is exactly 22 nucleotides long and have a guanine at their 5’.

#### RppH and TAP sample correlation

To determine the similarity between libraries, two approaches were used: (1) the genome was divided into equally sized bins and reads mapping to each bin were counted (DeepTools, multiBamSummary bins); or (2) reads mapping to a set of published 22G small RNA targets (DeepTools, multiBamSummary BED-file). Using either of those counts, the correlation coefficient (spearman and Pearson) between samples was calculated and plotted (DeepTools, plotCorrelation).

#### Differential gene expression

We used featureCounts (v1.4.6) with options -s 2 --ignoreDup to count the number of reads mapping to the annotated *C. elegans* genes (Ensembl, WBcel235). Read counts were then used in DESeq2 to determine differential expression between RppH and TAP treatments. Since the goal was to determine the experimental (technical) differences between both treatments, we considered each technical replicate as a biological replicate to perform the analysis.

#### 22G-RNA target annotation

The list of genes targeted by 22G-RNAs was compiled from previous publications (Claycomb et al., 2009; Conine et al., 2010, 2013; Gu et al., 2009; Phillips et al., 2014; Vasale et al., 2010). We then used the gene IDs to retrieve genomic locations from Ensembl using the bioconductor package biomaRt (Durinck et al., 2009).

## Results and Discussion

To address if RppH treatment enriches for 22G-RNAs as efficiently as TAP, we sequenced small RNAs of wild-type adult worms with prior treatment with TAP or RppH. To properly compare both treatments, we sequenced three technical replicates of TAP treatment and three technical replicates of RppH treatment, all derived from a unique biological sample.

We first determined if RppH treatment would affect the quality of the sequenced reads. QC reports generated by FastQC shows that the per base sequence quality of RppH-treated libraries is at least as good as that of TAP-treated libraries (**Supplementary Figure 1**). Other key indicators such as the average quality per read and sequence duplication levels are also similar in libraries treated with either method. Importantly, the three RppH-treated technical replicates show that these libraries are consistently of high quality.

The goal of both Rpph- and TAP-treatment is to enrich the library for 22G-RNAs. This class of small RNAs has two features that can be readily used to determine if and how successful the enrichment was: (i) most sequences are 22 nucleotides in length and; (ii) high proportion of sequences with a Guanine at their 5’ end. After adapter and UMI removal we plotted the distribution of read length in the sequenced libraries (**Figure 1A**). Reads were normalized to total number of sequenced reads. For comparison purposes, we added samples that were neither treated with TAP nor RppH, to set the baseline for enrichment. Untreated libraries have two abundant populations of reads with lengths of 21 and 23 nucleotides, accounting for 21U-RNAs and miRNAs, respectively, whereas in the RppH- and TAP-treated libraries there is a clear enrichment of reads with 22 nucleotides. Furthermore, in sequences originating from untreated libraries most reads contain a 5’ Uracil, but approximately 50% of reads from RppH and TAP-treated libraries have a Guanine at this position (**Figure 1B**). This shows that RppH-treatment is as effective as TAP in enriching for reads with 22G-RNAs.

**Figure 1.**
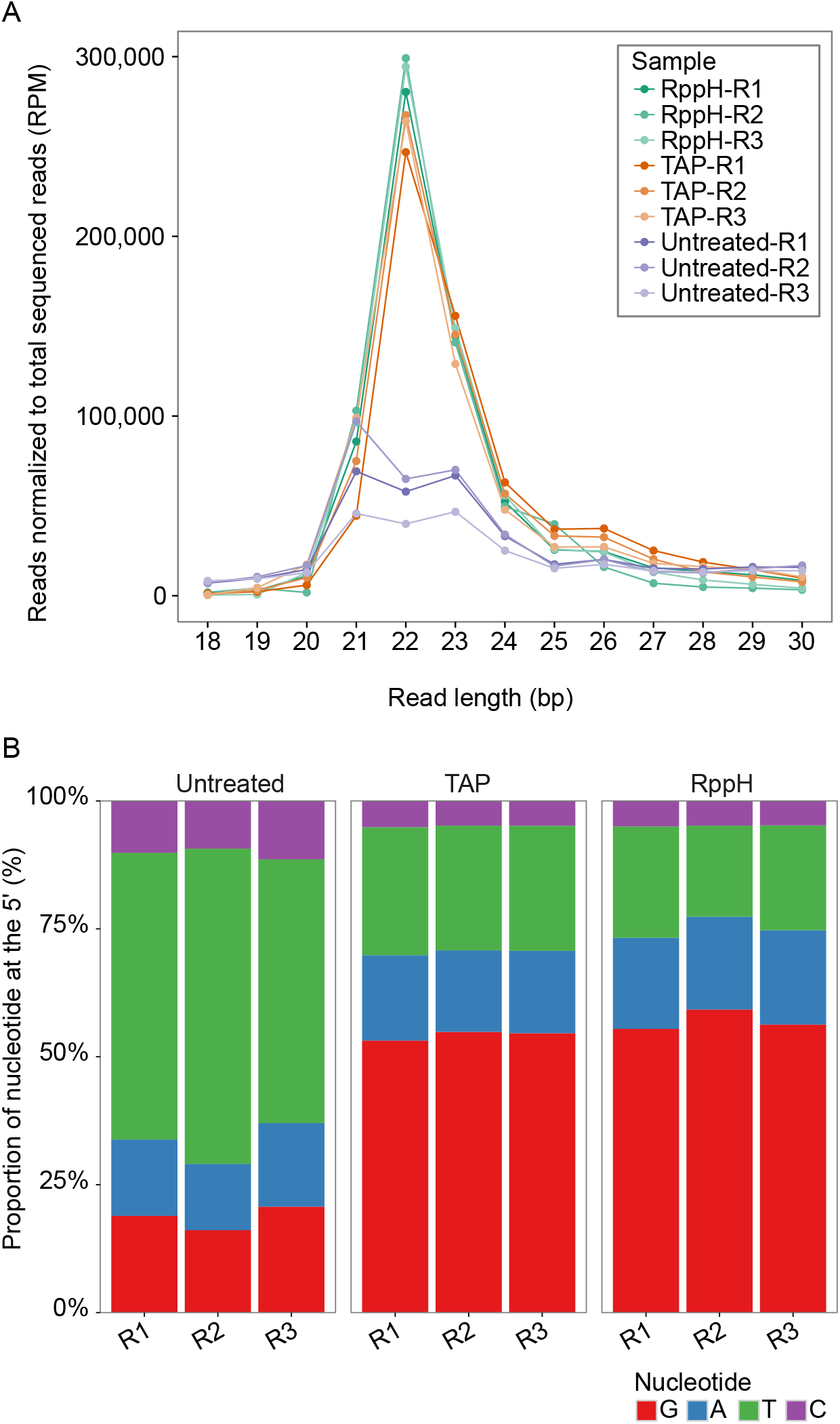
RppH-treatment enriches for sequences of 22 nucleotides in length and with 5’ guanine bias. (A) Distribution of sequence lengths normalized to the total number of sequenced reads for each library. RppH- and TAP-treated libraries show a larger number of 22 nucleotide long reads. (B) 5’ nucleotide bias in sequenced reads. R1-R3: Replicates 1-3.

For both treatments, reads were mapped to the *C. elegans* genome with equal efficiency, and similarly to untreated libraries (**Figure 2A**). Despite achieving a high enrichment of 22G-RNA sequences (**Figure 1**), it is possible that the TAP- and RppH-treatment select for different subpopulations of these small RNAs. To exclude this possibility, we selected, in the RppH- and TAP-treated libraries, only reads that are 22 nucleotides in length and have a guanine at the 5’ (22G reads), and tested if these reads map to the same genomic locations. All the follow-up analysis in this manuscript is done using these 22G-RNA reads. The overall 22G-RNA genomic coverage appears to be similar between RppH and TAP-treated libraries (**Figure 2B**, **Supplementary Figure 2**). To confirm this observation the genome was split in bins of equal lengths, we counted the number of reads mapped to genomic bins, and compared these for all samples (**Figure 2C**). The mapped positions of 22G-RNAs from TAP- and RppH-treated libraries are highly correlated (Spearman’s correlation > 0.86), and as expected are very different from those of the untreated libraries (Spearman’s correlation < 0.43). Technical replicates of the RppH treatment are also consistently highly correlated, indicating that the RppH treatment is reproducible.

**Figure 2.**
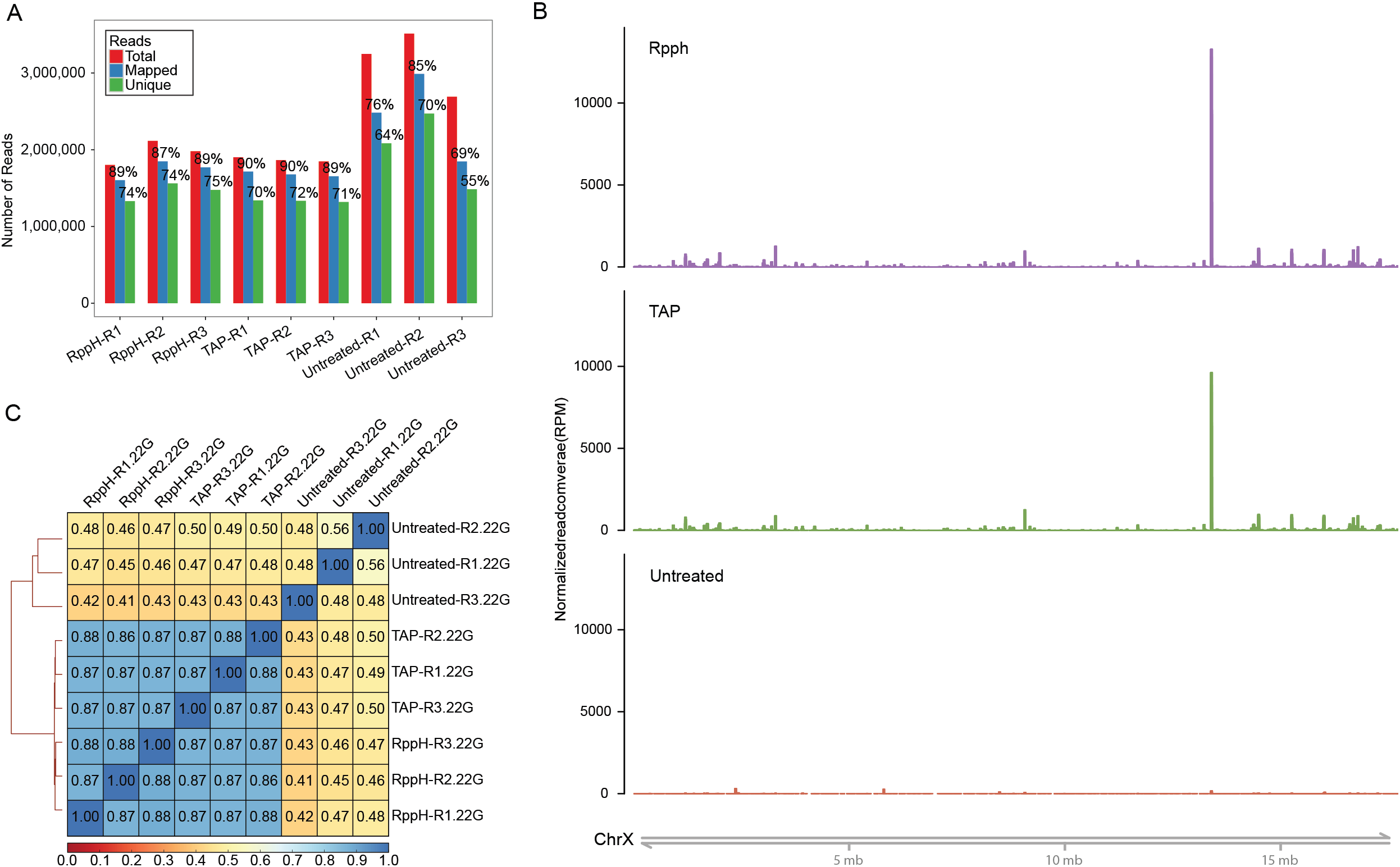
RppH-treatment does not affect the amount of reads mapped to the genome or the location of 22G-RNA reads, compared to TAP-treatment. (A) Efficiency of read alignment to the *C. elegans* genome. Total number of reads (red bars) refers to the number of reads that passed the preprocessing and quality filtering steps. Mapped reads (blue bars) are those that map at least once to a region in the genome, and unique reads (green bars) are those that align to a single genomic location. (B) Overview of normalized read coverage in chromosome X. Reads of 22 nucleotides in length and a guanine at the 5’ were isolated (22G) and their abundance plotted. For visualization purposes the three replicates of the libraries of each preparation method where merged. Plots for all chromosomes with individual replicates are provided in **Supplementary Figure 2**. (C) Spearman correlation matrix of all replicates obtained from 22G read counts to genomic bins. R1-R3: Replicates 1-3.

Even though the overall genomic location of RNA recovered from RppH or TAP treatment is very similar (**Figure 2B**, **Supplementary Figure 2**), there might be subtle differences between the treatments, possibly in genes known to be targeted by 22G-RNAs. To exclude this possibility, we determined if the RppH treatment recovered small 22G-RNAs that map to known 22G-RNA targets. We counted the number of reads mapping to lists of published target genes of certain small RNA pathway factors (Claycomb et al., 2009; Conine et al., 2010, 2013; Gu et al., 2009; Phillips et al., 2014; Vasale et al., 2010) and plotted sample correlations (**Figure 3A**). Replicates from TAP- and RppH-treated libraries are highly correlated (Spearman’s correlation >= 0.82), whereas untreated samples show very different patterns, (Spearman’s correlation <= 0.48 versus TAP or RppH). Furthermore, RppH and TAP libraries are enriched in 22G small RNAs in these regions, compared to untreated libraries (**Supplementary Figure 3**). We have also looked in detail into the coverage of known 22G-RNA targets, and confirmed that the level of genic reads resulting from either treatment show the same patterns (**Figure 3B**). This strongly suggests that RppH and TAP treatment enriches for 22G-RNAs that target the same genes. To further determine if there is some bias in either treatment, we performed differential gene expression analysis comparing TAP vs RppH and found only 1 gene differentially enriched between TAP and RppH libraries (F38E11.21, a non-coding RNA, FDR < 0.1). Importantly, F38E11.21 is not a known 22G-RNA target. From this, we conclude that RppH treatment faithfully recapitulates TAP treatment.

**Figure 3.**
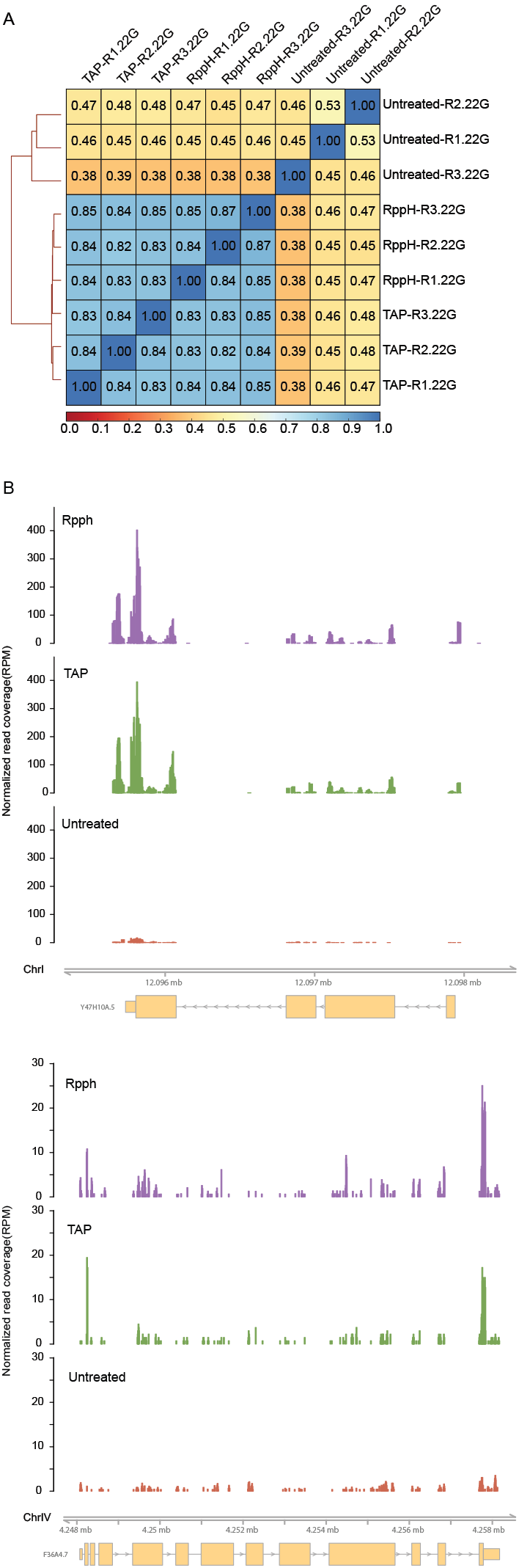
22G-RNA reads isolated with RppH- or TAP-treatment mapped similarly to known 22G-RNA targets. (A) Spearman correlation of reads mapping to known 22G-RNA targets. (B) Normalized read coverage at two known 22G-RNA targets, *Y47H10A.5* and *ama-1* (Corrêa et al., 2010; Gu et al., 2009). For visualization purposes the three replicates of each library preparation method where collapsed.

The unavailability of TAP from commercial sources eliminated a common tool used in the field of small RNA research, where, amongst other uses, TAP was used to enrich samples for 22G-RNAs in *C. elegans*. Whilst no consensus alternative for TAP has been found, others have used other alternatives to TAP and RppH (Houri-Ze’evi et al., 2016; Sapetschnig et al., 2015), however, no systematic comparison of these enzymatic treatments was performed. Here, we show that another decapping enzyme, RppH, is a suitable alternative to TAP. Small RNA-seq libraries of samples treated with RppH not only have similar number of 22G-RNA reads, but these also originate from the same genomic locations as that of TAP-treated libraries. While preparing this manuscript, we sequenced small RNAs of various other RppH-treated libraries (data not shown). Analysis of those small RNA sequencing datasets attested for the reproducibility of RppH treatment (data not shown). Overall, we can recommend RppH as a direct replacement of TAP as a small RNA pyrophosphatase to enrich RNA samples for 22G-RNAs with almost no changes to existing protocols or added costs.

---

**Figure S1.**
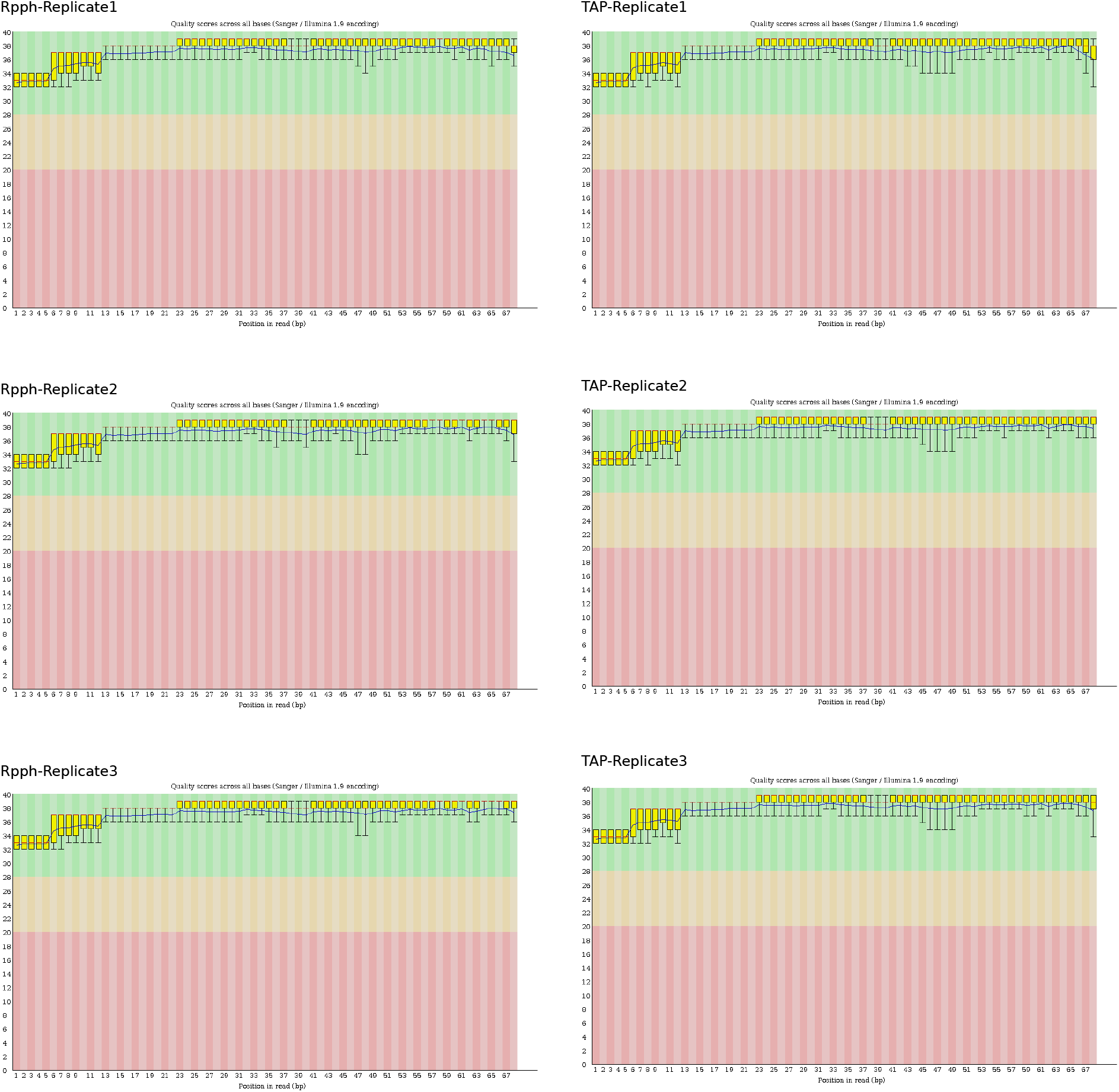
Per base sequence quality of TAP- and RppH-treated libraries.

**Figure S2.**
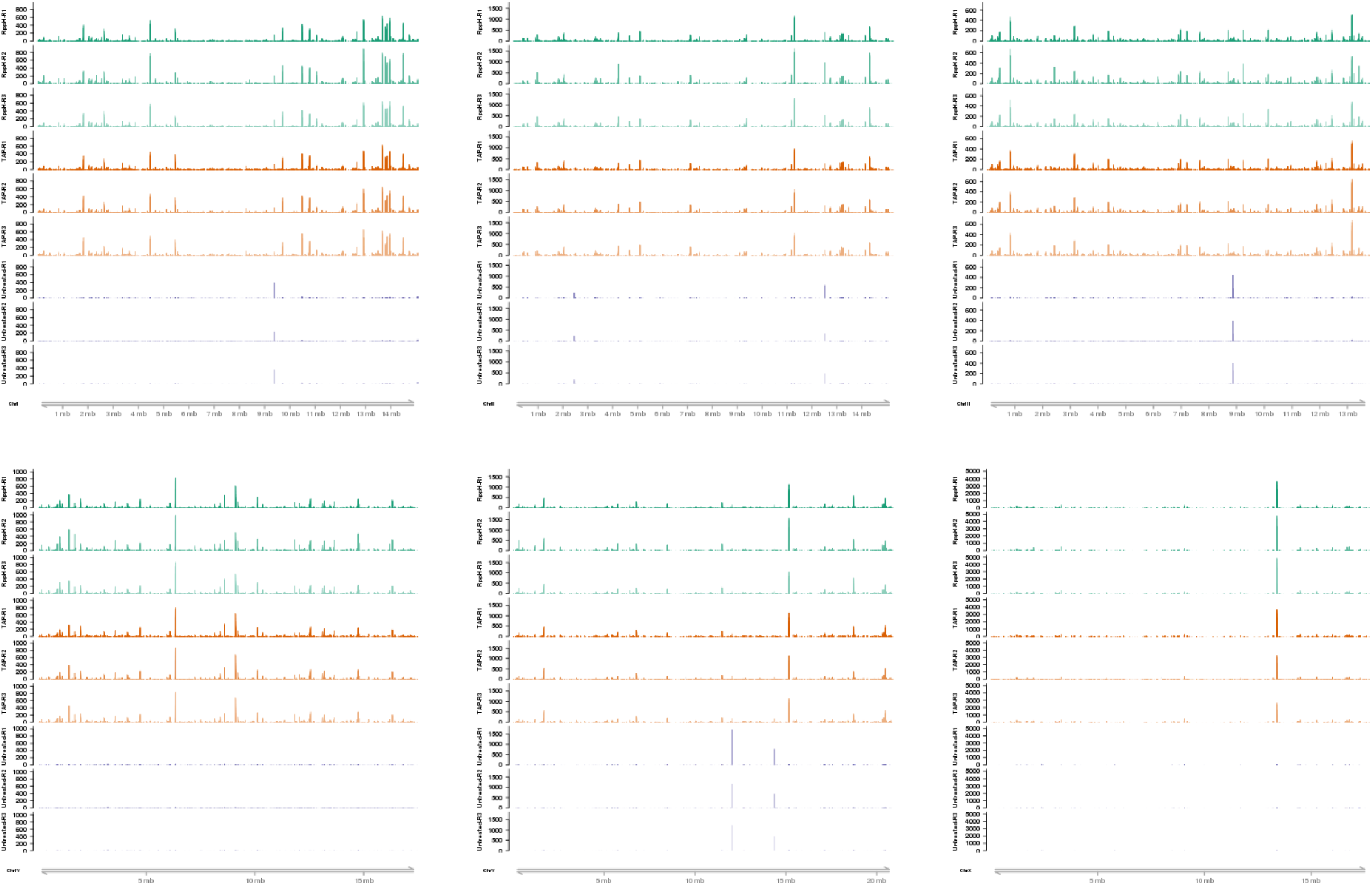
Genomic coverage of 22G-RNAs in TAP- and RppH-treated libraries. All chromosomes are shown.

**Figure S3.**
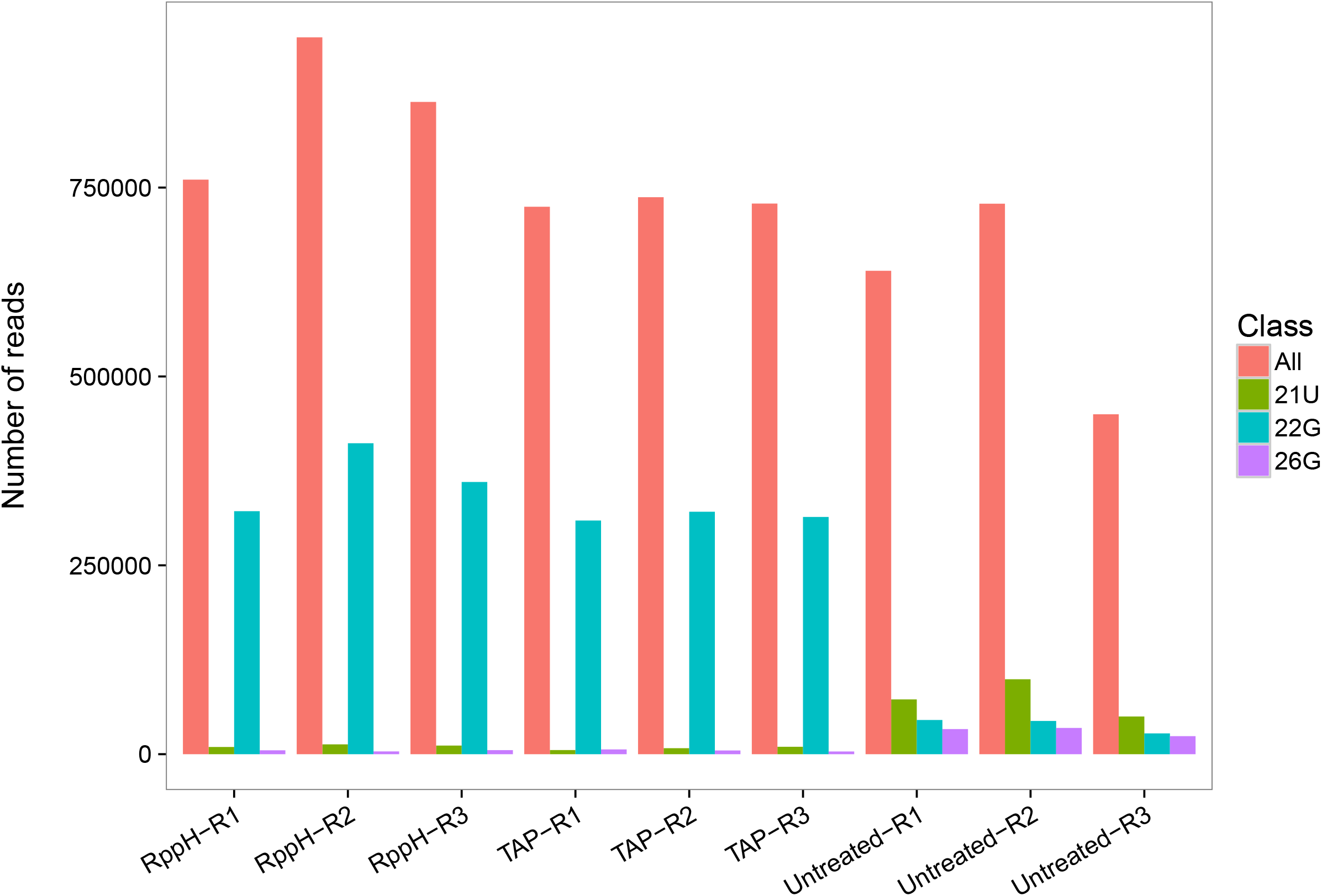
22G-RNA enrichment in 22G-RNA targets, as defined in other studies.

## Acknowledgements

We are grateful to Maria Placentino for comments on the manuscript.

## Author Contributions

Conceived and designed the experiments: MVA, AD, HL, MML. Performed the experiments: MVA, HL, MML. Analyzed the data: AD, MVA, RFK. Wrote the paper: AD, MVA, RFK.

## Data Availability

Sequencing data have been deposited to the NCBI Gene Expression Omnibus (GEO): GSE111995 for TAP- and RppH-treated samples; GSE103432 for the untreated samples, also see (Almeida et al., 2018).

